# Validation of Flavivirus Infectious Clones Carrying Fluorescent Markers for Antiviral Drug Screening and Replication Studies

**DOI:** 10.1101/2023.04.03.535319

**Authors:** Liubov Cherkashchenko, Nathalie Gros, Alice Trausch, Aymeric Neyret, Mathilde Hénaut, Gregor Dubois, Matthieu Villeneuve, Christine Chable-Bessia, Sébastien Lyonnais, Andres Merits, Delphine Muriaux

**Author notes:** These first authors contributed equally to this article (Liubov Cherkashchenko, Nathalie Gros, Alice Trausch). These senior authors were co–principal investigators (Andres Merits, Delphine Muriaux).

## Abstract

Flaviviruses have emerged as major arthropod-transmitted pathogens and represent an increasing public health problem worldwide. High-throughput screening can be facilitated by the use of viruses that express easily detectable marker proteins. Developing molecular tools such as reporter-carrying versions of flaviviruses for studying viral replication and screening of antiviral compounds therefore represents a top priority. However, the engineering of flaviviruses carrying either fluorescent or luminescent reporters remains challenging due to the genetic instability caused by marker insertion; therefore, new approaches to overcome these limitations are needed. Here, we describe reverse genetic methods which includes design and validation of infectious clones of Zika, Kunjin and Dengue viruses harboring different reporter genes for infection, rescue, imaging and morphology using super-resolution microscopy. It was observed that for different flaviviruses constructs with identical design displayed strikingly different genetic stability while corresponding virions resembled wild-type virus particles in shape and size. A successful strategy was assessed to increase stability of rescued reporter virus and permit antiviral drug screening based on quantitative automated fluorescence microscopy and replication studies.

## INTRODUCTION

The *Flaviviridae* family includes four genera: *Flavivirus*, *Pestivirus*, *Pegivirus* and *Hepacivirus* (https://ictv.global/taxonomy). Genus *Flavivirus* includes several arthropod-borne viruses usually infect insects but some can be responsible for a significant number of human diseases primarily caused by dengue virus (DENV), Zika virus (ZIKV), West Nile virus (WNV), yellow fever virus (YFV), Japanese encephalitis virus (JEV), and tick-borne encephalitis virus (TBEV). In the majority of the cases, transmission occurs horizontally between mosquitoes (often *Aedes* or *Culex*) or ticks and vertebrate hosts (1). It is worth of noting that flaviviruses represent a group of pathogens that are dangerous not only for humans but also for different mammalians (including domestic species); the latter often serving as reservoirs and thus contributing to the adaptation of viruses to a new environmental conditions and more efficient transmission to humans (2). In this study, we focused on investigation of three members of genus *Flavivirus*: DENV (genotypes 2 and 4), ZIKV and Kunjin virus (KUNV), an Australian subtype of WNV.

The flavivirus genome is a positive-strand RNA of about 11,000 bases containing a single open reading frame (ORF) that is flanked by two untranslated regions (UTRs) located at the 5’ and 3’ ends of virus genome. Flavivirus genome has cap structure located at the 5’ end but lacks the 3’ poly(A) tail. The absence of poly(A) is compensated for by the interaction of 3’UTR with poly(A) binding protein (PABP). Flavivirus ORF encodes for a polyprotein precursor of viral proteins that is cleaved by viral and host proteases into three structural proteins (C (capsid), prM(M) (membrane), and E (envelope)) and seven non-structural (NS) proteins (NS1, NS2A, NS2B, NS3, NS4A, NS4B, and NS5). The NS proteins play multiple roles in infection cycle including in viral RNA replication, assembly of viral particles and evasion of host immune response. The structural proteins are involved in the formation and release of viral particles (3, 4).

The clinical manifestations of flavivirus infection range from mild (often asymptomatic) infection to infection associated with severe symptoms (5). Such a divergence is mostly caused by different tropism of viruses or/and their ability to counteract the host immune response. There are approximately 400 million flavivirus infection every year, large majority caused by four genotypes of DENV (DENV1-4) making it one of the most medically important causative agents of human diseases (6). The clinical presentation of DENV infection includes development of disease - dengue fever - and also more severe diseases - dengue hemorrhagic fever and dengue shock syndrome (7). There are around 100 million symptomatic DENV cases each year and about 40 000 of these lead to the development of severe illness (8). In contrast to DENV, symptomatic infection by ZIKV is mostly associated with mild symptoms such as headache, joint pain or cutaneous rash. In some cases, ZIKV infection can progress into Guillain-Barré syndrome accompanied with progressive muscular paralysis in 20-30% of patients (9, 10). Moreover, retrospective studies on the past outbreaks (Yap island, French Polynesia and Brazil) established a causality between ZIKV infection and microcephaly in newborns in women infected during pregnancy (11).

Natural hosts of DENV and ZIKV are primates and these viruses are capable for establishment urban (human-mosquito-human) transmission cycle. Interestingly, WNV has wider range of vertebrate host including birds serving as natural reservoirs; for this virus horses and humans represent dead-end hosts. WNV comprises at least 7 genetic lineages. KUNV belongs to the lineage 1b of WNV and is endemic to Australia. Infection with KUNV mostly lead to the development of mild symptoms in humans but the infection is lethal for horses (12). Currently, there are no specific treatments for flavivirus infections as well as efficient methods to control the spread of their arthropod vectors. Vaccines capable to prevent infection are available for YFV, JEV and TBEV. The situation with DENV is more difficult as it comprises four antigenically different serotypes. Development of vaccine for DENV has therefore been hampered by the existence of cross-reaction between antibodies against one serotype with viruses belonging to another serotype; in such case virus may escape neutralization which leads to an antibody dependent enhancement (ADE) believed to be responsible for establishment of severe forms of the DENV disease (13).

Antiviral drug discovery requires robust screening and antiviral activity assays. Development of such assays for different flaviviruses has therefore been in the focus of numerous studies. Standard screening assays for antivirals employ cytopathic effect (CPE) based readouts (14, 15, 16), reverse transcription quantitative polymerase chain reaction (RT-qPCR) based analysis of viral nucleic acids (17, 18, 19) or immunofluorescence staining of virus-encoded proteins (20, 21). All of these methods have been successfully applied to quantify the effect of antivirals on flavivirus infection; however, they remain time-consuming and expensive. Therefore, development of new approaches using reverse genetics of flaviviruses is necessary. Generation of infectious clones and recombinant flaviviruses harboring marker genes (encoding for fluorescent or luminescent reporters) allows easier tracking the course of the infection, quantification of the efficiency of antiviral treatments and consequently to screen and verify the antiviral drugs or to identify the host factors involved in the flavivirus infection. In the context of the study, DENV, ZIKV and KUNV reporter viruses have been designed and their properties were described; this allowed the development of efficient bioluminescence or image-based high-throughput assays applicable to drug discovery (22, 23, 24, 25, 26, 27). The caveat with the use of this approach is that the reporter viruses should have properties (including replication efficiency) similar to wild type (wt) viruses, and, furthermore, they have to be genetically stable. In practice, production of clone-derived recombinant flaviviruses corresponding to these requirements, has been challenging. The main reasons for this are low tolerance of flaviviruses to insertions and attenuation (reduced replication) of reporter viruses. To cope with these issues, here, following steps were previously applied: (i) improving reverse genetic systems for ZIKV, KUNV, DENV2 and DENV4 in order to allow easy and efficient manipulation of the corresponding infectious clones (23), (ii) stabilization of the reporter expression cassette by duplication of the capsid region (23, 27, 28). Nevertheless, genetic stability and durable reporter expression during long-term passage in tissue culture remained a key issue (23, 26, 28).

In the context of the current study, we aimed to develop and test newly designed wt infectious clones of KUNV, DENV2 and DENV4 as well as their versions harboring NanoLuciferase (NLuc), mCherry and oxGFP reporters constructed based on a strategy previously successfully applied to ZIKV (29). Herein, we present the data related to the strategies used for the construction of infectious clones and assessment of the growth kinetics of viruses rescued from these clones. Genetic stability of the marker-coding viruses was analyzed by monitoring the fluorescence or luminescence signals from the cells infected with the corresponding viruses. We also characterized the structures of the corresponding virions, variations in their sizes and shapes as well as their distribution in cellular compartments using transmission electron microscopy (TEM) on fixed infected cells and atomic force microscopy (AFM) on live viruses (57). Finally, we validated the use of obtained recombinant viruses in fluorescent or luminescent-based drug screening assays by testing NITD008 as a reference molecule harboring pan-flavivirus activity.

## RESULTS

### Design and construction of icDNA clones of DENV2, DENV4 and KUNV

Rescue of the virus from the infectious cDNA (icDNA) clone of an RNA virus enables various modification and/or manipulation of the RNA genome. Development of reverse genetic systems for different RNA viruses has been advanced over the last decades. The RNA transcripts derived from cDNAs of positive-strand RNA viruses are considered to be infectious i.e. their transfection into susceptible cells results in replication and successful recovering of the infectious viruses (31). For most of positive-strand RNA virus families this approach is relatively straightforward; however, for some of viruses, the development and use of reverse genetics is more challenging. Flaviviruses represent an example of the latter. It has been shown that some sequences from flavivirus genome encode proteins with high level of cytotoxicity for *E. coli*, a bacterium most commonly used to propagate plasmids containing viral icDNAs (32). The expression of such proteins via activity of cryptic bacterial promoters found in the cDNAs of flaviviruses is therefore harmful for the bacteria harboring corresponding plasmid leading to counterselection resulting in instability of icDNA plasmids.

Here we aimed construction of the icDNAs based on the NCBI sequences for DENV2 (GenBank: U87411.1), DENV4 (GenBank: AF326573.1) and KUNV (GenBank: AY274504) using strategy previously described for ZIKV (GenBank database: KJ776791) (33). Briefly, the SP6 promoter was placed upstream of the sequence corresponding to the 5’ end of virus genome while a cleavage site of restriction endonuclease was placed immediately downstream of sequence corresponding to the 3’ end of the genome. This allowed performing run-off *in vitro* transcription to obtain transcripts corresponding to the viral genome RNA that could be used for the efficient rescue of the infectious viruses. The sequences corresponding to virus genomes were assembled from synthetic DNA fragments; assembly was performed in a single copy plasmid to ensure efficient propagation of cloned cDNAs in bacterial cultures and to prevent possible re-arrangements in the cDNAs of flaviviruses (34). No instability issues were observed during plasmids construction and propagation of plasmids containing cDNAs of KUNV, DENV2 and DENV4. Of note, this approach failed with icDNA of DENV3 probably indicating extreme toxicity of the latter plasmid for *E. coli*.

To extend the study, along with preparation of the icDNA clones containing the wt sequence of DENV2, DENV4 and KUNV, icDNAs corresponding to recombinant viruses with inserted reporter-encoding genes were also constructed. The reporters allowing detection of the infection by either quantification of the marker expression/activity (NanoLuciferase, NLuc) or visually due to the fluorescence of the marker (oxGFP, mCherry) were used. It has been established that the position and insertion strategy of sequence encoding for marker in flavivirus genome plays a crucial role in the genetic stability of recombinant virus and affects the speed of loss of marker over passages (35). In our study the design previously used for construction of stable reporter viruses of ZIKV was applied for DENV2, DENV4, and KUNV. In this design the sequence encoding marker protein is placed between native sequence encoding for flavivirus capsid protein and that of the Foot-and-Mouth disease virus (FMDV) 2A autoprotease followed by codon-altered copy of sequence encoding for the capsid protein (Figure 1A). Thus, upon translation of recombinant genome, the marker is released from the first copy of capsid protein by mechanisms naturally used for release of flavivirus capsid from the polyprotein while FMDV 2A cleaves itself from the following copy of capsid protein allowing efficient release of reporter with no or minimal disturbance of maturation of flavivirus structural proteins as was confirmed by immunoblots (Figure S1A).

**Figure 1.**
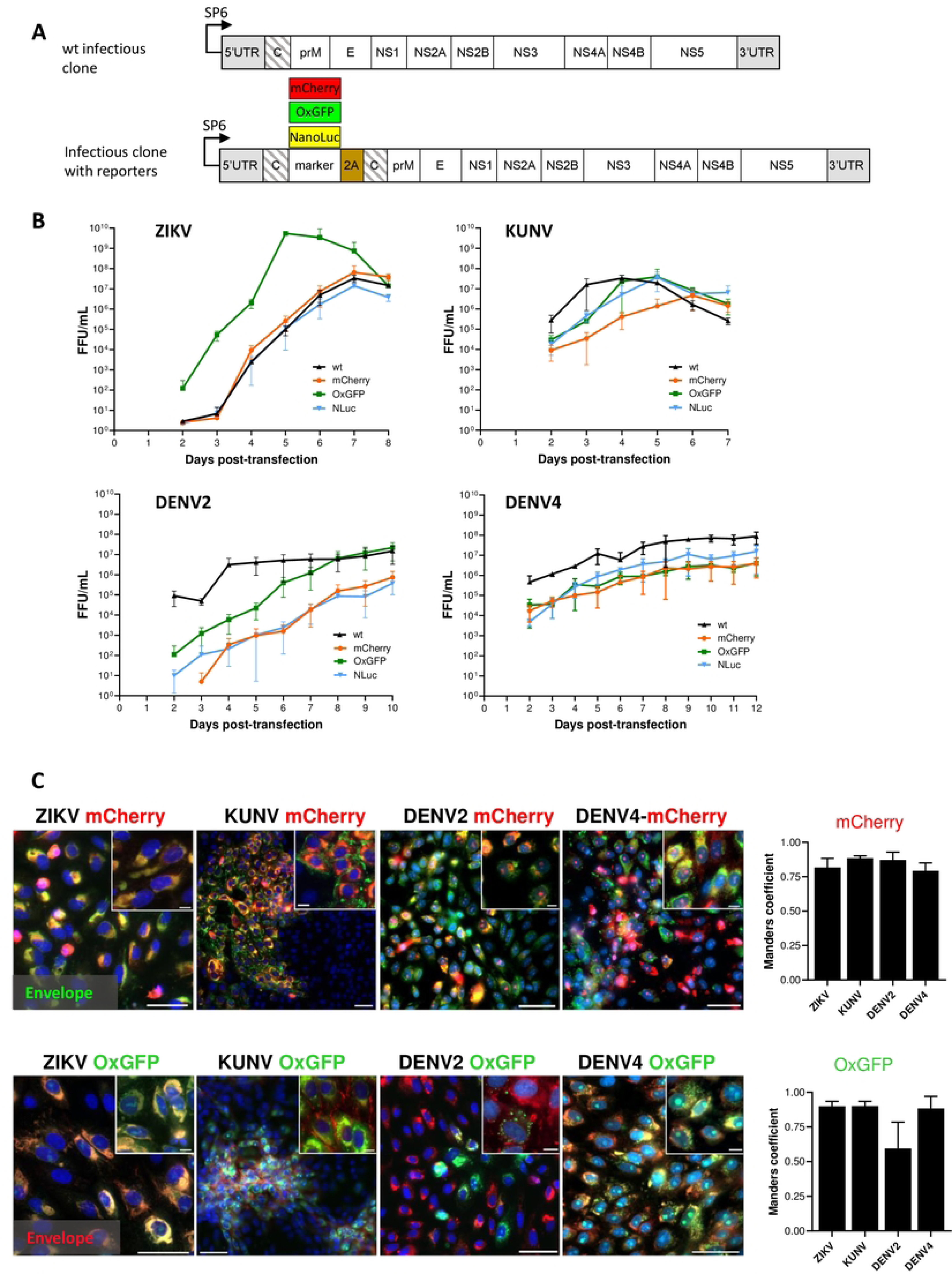
Design of wt and reporter containing icDNA of flaviviruses. (**A**) Schematic presentation of the viral constructs used in the study. The reporter (mCherry, oxGFP, NLuc) and Foot-and-Mouth-Disease virus 2A sequence (2A) were placed at the junction of two full-length capsid sequences (shown with slanted lines). The second sequence was codon-altered using synonymous substitutions to avoid the homologous recombination. (**B**) Rescue and growth kinetics of recombinant ZIKV, KUNV, DENV2 and DENV4 variants. After transfection of the *in vitro* transcribed RNA, the supernatant was collected at indicated time points, and the viral titer was quantified by FFA. (**C**) Left panel: localization of fluorescent reporter markers and envelope proteins. Vero cells were infected with P_0_ stocks of ZIKV-mCherry, ZIKV-oxGFP at MOI 0.1, and with P_0_ stocks of DENV2-oxGFP, DENV2-mCherry, DENV4-mCherry, DENV4-oxGFP, KUNV-oxGFP and KUNV-mCherry at MOI 0.05. At 72 h post infection cells were fixed and stained against viral enveloppe protein (green - for cells infected with viruses expressing mCherry and red - for cells infected with viruses expressing oxGFP). mCherry and oxGFP signals were detected by confocal microscopy; nuclei were counterstained with Hoechst (blue). Scale bars: 10 µm. Right panels: colocalization of red and green fluorescence (Manders’ coefficient) for ZIKV-mCherry (1082 cells), KUNV-mCherry (680 cells), DENV2-mCherry (642 cells), DENV4-mCherry (600 cells), ZIKV-oxGFP (716 cells), KUNV-oxGFP (622 cells), DENV2-oxGFP (706 cells) and DENV4-oxGFP (645 cells). Data are represented as mean +/-SD from three independent experiments.

### Rescue and properties of wt and reporter-expressing variants of DENV2, DENV4, ZIKV and KUNV

Rescue of DENV2-wt, DENV4-wt, ZIKV-wt, KUNV-wt as well as their variants encoding mCherry, oxGFP or NLuc reporters was performed in Vero cells (Figure 1). Focus forming assay (FFA) using pan-flavivirus E protein-specific mouse MAb (4G2) was used to determine the titers of each of the rescued viruses at different days post transfection (dpt). Supernatants designated as P_0_ stocks were collected at the peak of virus release and used to infect new cells for imaging and further analysis. It allowed to identify the percentage of the infected, marker expressing cells, in the total pool of infected enveloppe-labelled cells (Figure 1C).

It was observed that ZIKV-wt, ZIKV-mCherry and ZIKV-NLuc were cytopathic and showed similar exponential growth. ZIKV-wt become detectable at 2 dpt and a maximum titer of 6.5 x10^7^ FFU/mL was reached at 7 dpt (Figure 1B). Curiously, but for unknown apparent reasons, at early time points ZIKV-oxGFP had titers that were approximately 20x higher than other clone-derived variants of ZIKV reaching 5.5×10^9^ FFU/mL at 5 dpt; however, after that the titer decreased and by 8 dpt and become comparable with the other ZIKV constructs used in the study (Figure 1B). After transfection with transcripts of ZIKV-mCherry, the first mCherry-expressing cells were observed at 4 dpt and their number increased in following days (Figure S2). Interestingly, despite a high viral titer, ZIKV-oxGFP infected cells showed a low fluorescent signal-to-noise ratio (SNR ∼10), which reduced the interest in ZIKV-oxGFP as a reporter virus. In contrast, ZIKV-mCherry infected cells showed a robust fluorescent SNR of ∼40, thus increasing the value of the use of this virus as a reporter in fluorescence-based experiments, for instance, drug screening.

Cells transfected with transcripts of icDNAs of KUNV displayed stronger cytopathic effects (CPE) than that observed for ZIKV: a complete CPE was observed already by 7 dpt. Coherently, rescue of KUNV was also more rapid: at 2 dpt the titer of KUNV-wt was already 2.8×10^5^ FFU/mL and reached maximum values of 3.4×10^7^ FFU/mL at 4 dpt (Figure 1B). Slight delay in the development of CPE was observed for KUNV-oxGFP and KUNV-NLuc which was reflected in reduced titers at 2 dpt. Up to 5 dpt, the titers of KUNV-mCherry were the lowest; however, at 6 dpt they reached the level similar to those of KUNV-wt, KUNV-oxGFP and KUNV-NLuc (Figure 1B). As an example, mCherry-reporter fluorescence were detected in viral clone-transfected cells at 4,5,7 dpt up to 12dpt depending on the virus (Figure S2).

Additionally, the replication of reporter viruses was confirmed by staining with viral envelope (E) antibodies (Figure 1C). Interestingly, fluorescent microscopy showed a distinguishable feature of recombinant KUNV replication: both mCherry and oxGFP signals were mostly detected in large intracytoplasmic structures in infected cells, reflecting the co-localization of capsids, whereas the localization of oxGFP signal in the nucleoli was observed in only a limited number of cells. Viral E proteins were associated with the plasma membrane in both cases (KUNV mCherry and oxGFP) (Figure 1C).

Rescue of DENV2 and DENV4 was considerably slower than that of KUNV. Interestingly, in contrast to ZIKV and KUNV, no decrease of viral titers was observed at late time points (up to 12 dpt); instead virus titers either reached a plateau level (as observed for DENV2-wt) or continued to increase slowly (Figure 1B). While DENV2-wt rapidly reached high titers (approximately 1.5×10^7^ FFU/mL), lower titers were observed for all of the reporter harboring variants of this virus. For DENV2-oxGFP this was observed for earlier time points and ultimately the virus reached titers similar to those of DENV2-wt. In contrast, for DENV2-mCherry and DENV2-NLuc lower titers were observed during all time course of the experiment and the maximal titer remained as low as 5×10^5^ FFU/mL (Figure 1B). Similar “behavior” was observed for reporter variants of DENV4 – for all of these (including DENV4-oxGFP) titers remained lower than that of DENV4-wt for all the course of experiment (Figure 1B). Similarly to the ZIKV-oxGFP, the fluorescent signal in DENV2-oxGFP and DENV4-oxGFP positive cells remained dim and was detected as foci (DENV2) or diffusely distributed in the cell cytoplasm and the nucleus (DENV4). For all recombinant viruses harboring mCherry or oxGFP markers viral E protein predominantly accumulated in the cell cytoplasm (Figure 1C) where it co-localized with fluorescent marker protein. For DENV4-mCherry and DENV4-oxGFP colocalization was at 87% and 88% (Manders’ coefficient value), the respectively; similar colocalization was also observed for recombinants based on ZIKV and KUNV (Figure 1C). For both of DENV2-mCherry and DENV4-mCherry infected cells, the fluorescent mCherry signal was distributed within the cell cytoplasm and in the nucleus (Figure 1C). The lowest co-localization for viral E protein and fluorescent marker was observed for DENV2-oxGFP. Most likely this is not due specific properties of the virus/marker combination as estimation of the percentage of the cells positive only for oxGFP in the total population of infected cells revealed a rapid loss or inactivation of oxGFP marker (Figure 1C)

Overall, the results of these experiments demonstrated efficient rescue of recombinant flaviviruses from the RNA transcripts and revealed dynamics of their replication allowing to distinguish specific phenotypic features of infection. In addition, all oxGFP and mCherry expressing constructs with the exception of DENV2-oxGFP, showed high level (Mander’s coefficient value >80%) co-localization coefficients between the viral E protein and their respective marker (Figure 1C). As expected, the replication of ZIKV-wt, KUNV-wt, DENV2-wt and DENV4-wt was (with notable exception of ZIKV-oxGFP that we could not explain reasonably) more robust than that of viruses containing genes encoding for fluorescent or NLuc marker. This data indicates that insertion of the marker sequences had an impact on the level of RNA replication and/or virion formation and release.

### Assessment of genetic stability of recombinant virus genomes in cell culture

Flaviviruses carrying reporters are known to face instability problems. Due to mutations and recombination events the reporter genes are often lost during serial passaging of these viruses. The stability of recombinant flavivirus genome depends on multiple factors including virus species, marker insertion strategy and the used marker gene (36). Once marker is lost the resulting virus will eventually overgrow the parental recombinant; the speed of this outcompeting depends from conditions of infection (MOI) and from differences in growth kinetics of competing viruses. All marker-containing viruses, except for ZIKV, analyzed in previous experiment replicated slower than their wt counterparts (Figure 1B) demonstrating growth advantage of wt virus and indicating that viruses that have lost a sequence encoding for marker very likely also have growth advantage. Therefore, we analyzed the marker stability of all above described viruses with reporters by performing four passages in Vero cells with following quantification of the infectious titer using FFA (immunostaining against E) and analysis of marker expression by fluorescence microscopy (mCherry), fluorescence microscopy with immunostaining (anti-GFP) or luminescence measurement (NLuc) (Figure 2A).

**Figure 2.**
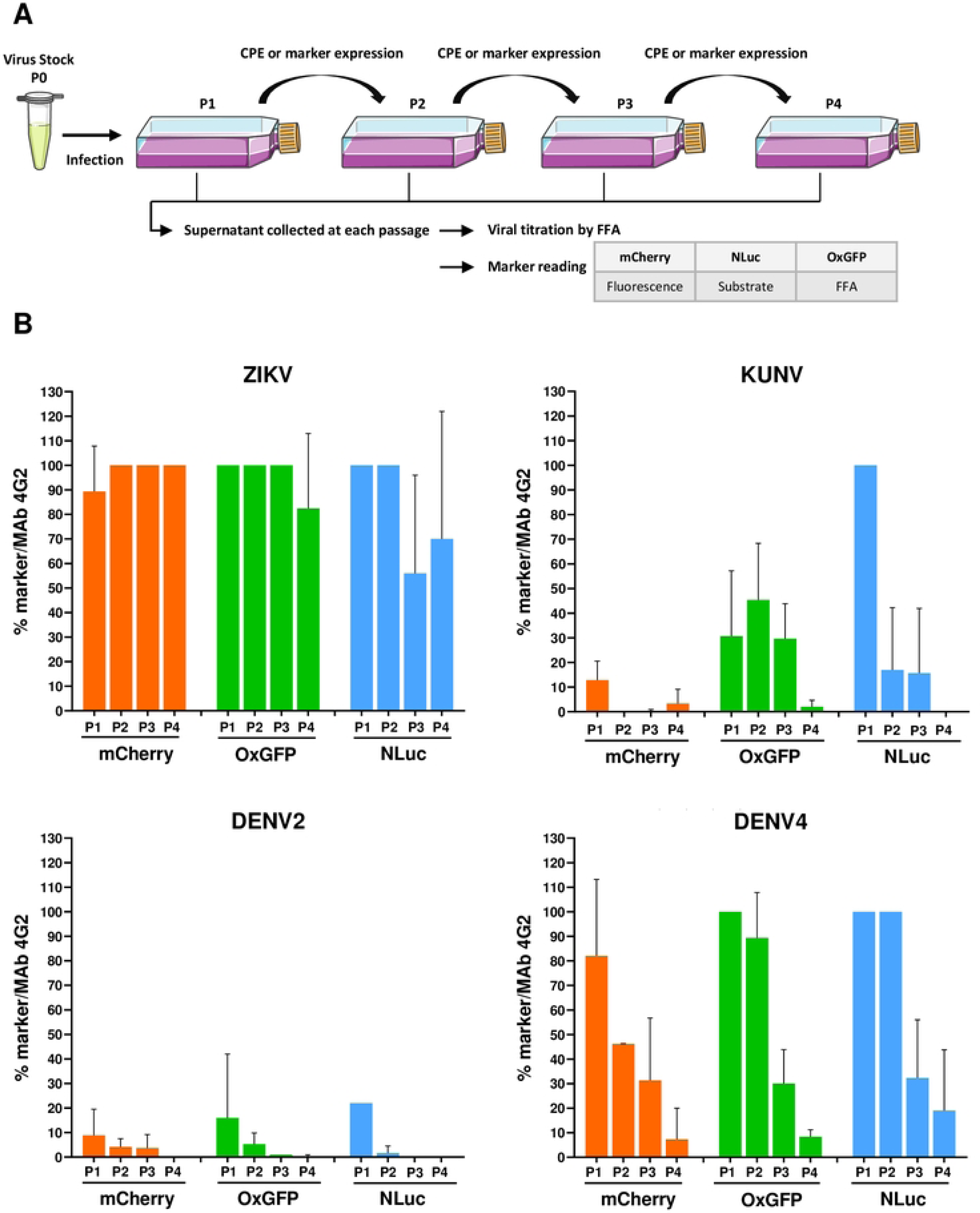
Evaluation of the genetic stability of -mCherry, -oxGFP- and -NLuc reporter carrying flaviviruses. (**A**) Schematic representation of the experimental protocol used to perform marker stability assay. Vero cells were infected with P_0_ stocks of recombinant ZIKV, KUNV, DENV2 or DENV4 with mCherry, oxGFP or NLuc markers and incubated until development of CPE or visible marker expression. P1 stocks were collected and used to infect Vero cells for the next passage until passage 4 (P1 to P4 stocks). Viral stocks were titrated by FFA using a mouse pan-flavivirus anti-Env antibody (Mab 4G2). mCherry signal was measured by quantification of fluorescence using a Cellomics ArrayScan VTI microscope. oxGFP expression was detected by immunofluorescence assay using an anti-GFP antibody. For the measurement of NanoLuciferase activity, a second 96-well plate was infected with P1 to P4 stocks according to the end-point titration method and NLuc was measured with the Nano-Glo Luciferase Assay System after cell lysis, on a microplate reader. (**B**) Evolution of the transgene expression through serial passages for each virus. The data represent the expression of the marker reported to the infectious titer. All experiments were performed in triplicate and results are represented as mean +/-SD.

Coherent with previous study (37) all recombinant variants of ZIKV displayed a relatively stable marker expression over four passages, with ≥ 80% cells infected with P_4_ stock of ZIKV-mCherry or ZIKV-oxGFP expressing corresponding marker (Figure 2B). Results obtained for ZIKV-NLuc displayed larger variation indicating partial loss of the inserted sequence during late passages; however, around 50% of cells infected with P_3_ or P_4_ stocks did express NLuc (Figure 2B) as measured by TCID50 using both FFA and direct NLuc measurement. In sharp contrast, loss of marker expression was already detected for P_1_ of KUNV-mCherry and KUNV-oxGFP; cells infected with P_2_ of these viruses revealed almost complete (mCherry) or >50% (oxGFP) loss of marker expression (monitor by FFA with a 4G2-antibody labelling). Marker expression (4G2 envelope) was detected in all cells infected by P_1_ of KUNV-NLuc; however, in subsequent passages the percentage of cells infected with the viruses expressing marker diminished and by passage 4 the marker expression was completely lost (Figure 2B). The effect was similar, albeit even more pronounced, for all recombinant DENV2 variants. Approximately 90% of loss of marker expression occurred already during the first passage and expression of markers was not observed during subsequent passaging (Figure 2B). Somewhat surprisingly, DENV4 reporter harboring variants were clearly more stable. mCherry expression was almost uniformly detected in cells infected with P_1_ of DENV4-mCherry followed by gradual decrease of marker positive infected cells over the next three passages. Both DENV4-oxGFP and-NLuc viruses were stable for two passages; however rapid loss of markers was observed for P_3_ and P_4_ stocks of these viruses (Figure 2B).

### DENV2-mCherry can be stabilized by truncation of the sequence encoding for the first copy of capsid protein

Instability of reporter-expressing recombinant flaviviruses has led to the development of various approaches aimed to overcome the problem. For viruses harboring marker between two copies of sequences encoding for capsid protein (Figure 1A) shortening of the first (native) copy of capsid protein gene to the length sufficient to preserve *cis-*active RNA elements located in this region has been successfully used; sometimes such a truncation is combined with additional modifications of the insertion region (38, 39). For some of flaviviruses the optimal length of the 5’ copy of capsid encoding sequence has been determined to be 35 or 38 codons; more extensive truncations have been shown to cause unpredictable recombination of the genome or loss of the marker (38, 40). To determine whether this approach can be used to stabilize highly unstable DENV2-mCherry, a deletion of the residues 39-114 in the first copy of capsid was made (Figure 3A). The rescue and properties of corresponding virus (designated as DENV2-Ct-mCherry) were compared with those of DENV2-mCherry (Figure 3B, C).

**Figure 3.**
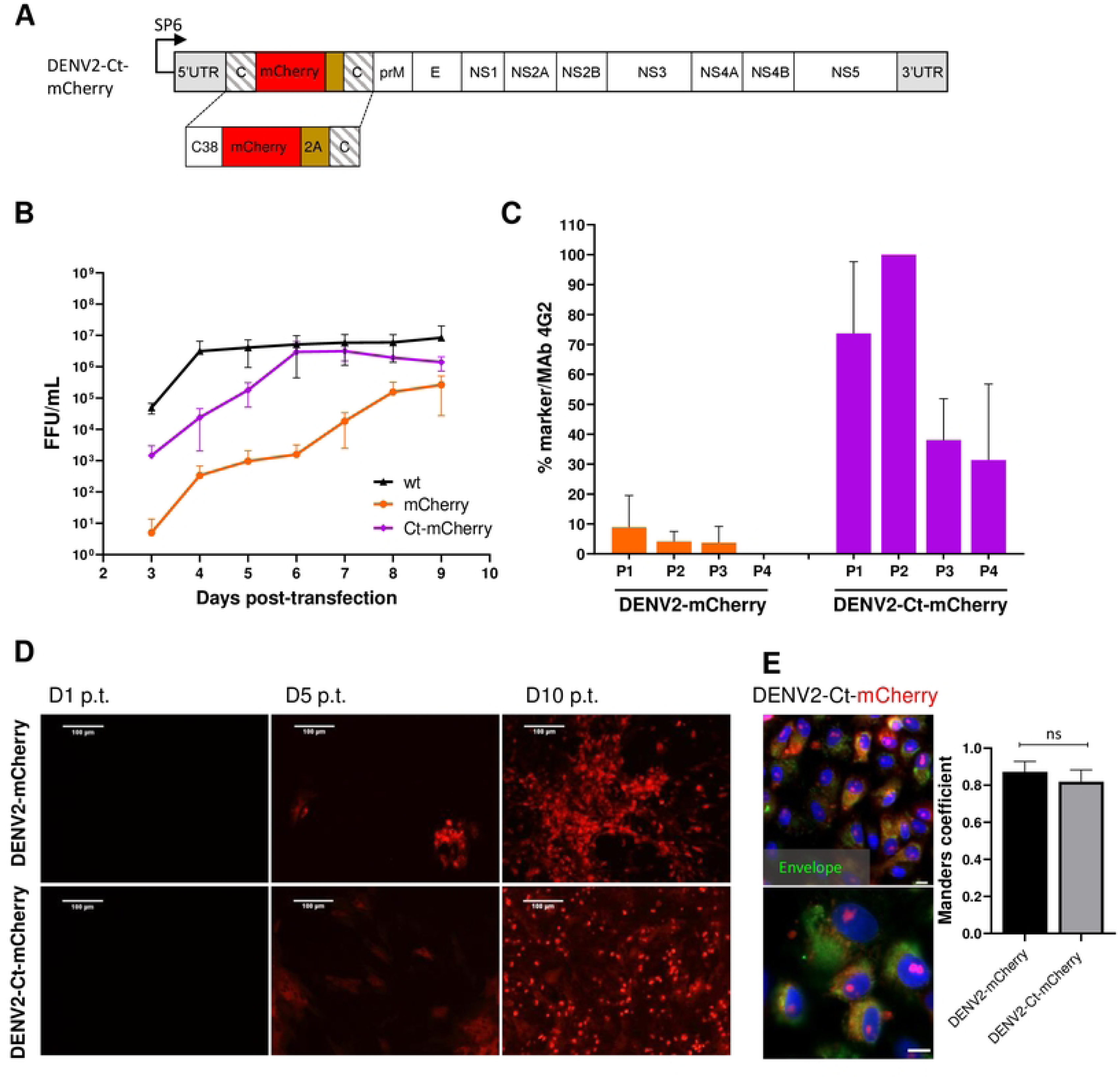
Comparison of the growth kinetics and genetic stability of DENV2-mCherry and DENV-Ct-mCherry. (**A**) Schematic representation of DENV2-Ct-mCherry icDNA clone. The first copy of the capsid protein was truncated by removing residues 39-114. The rest of the design corresponds to the icDNA clones of viruses with full-length capsid duplication. (**B**) Growth kinetics of recombinant DENV2-wt, DENV2-mCherry and DENV2-Ct-mCherry. (**C**) Comparison of marker stability of recombinant DENV2-mCherry and DENV2-Ct-mCherry over four passages. All experiments were performed in triplicate and results are represented as mean +/-SD. (**D**) Evaluation of replication dynamic DENV2-mCherry or DENV2-Ct-mCherry based on the increasing intensity of mCherry signal in the virus-infected cells over indicated time. (**E**) Left panel: confocal images of mCherry fluorescence (red) and DENV2 envelope protein (green) at 72h p.i. at MOI 0.05 (scale bars: 50 µm and 10 µm respectively). Right panel: colocalization of red and green fluorescent signals (Manders’ coefficient) for DENV2-mCherry (642 cells) and DENV2-Ct-mCherry (616 cells). Data represent mean +/– standard error of mean, n = 3, “ns” stands for “not significant”. Analysis of variances (ANOVA) was performed followed by a Tukey’s multiple comparisons test with GraphPad Prism 9.0.

Compared to DENV2-mCherry, higher titers were obtained for DENV2-Ct-mCherry over the time course of the virus rescue experiment. By 6 dpt the titers of DENV2-Ct-mCherry reached a plateau level (around 5×10^6^ FFU/mL) that was close to that of the DENV2-wt (Figure 3B). Remarkably improved genetic stability was also observed: all or nearly all cells infected with P_1_ or P_2_ stocks of DENV2-Ct-mCherry were positive for mCherry expression; in cultures infected with P_3_ or P_4_ stocks the loss of marker became more evident yet still 30 to 40% of infected cells were positive for mCherry (Figure 3C). No clear difference in the time scale of detection of mCherry fluorescence in reporter-positive cells was observed between DENV2-mCherry and DENV2-Ct-mCherry (Figure 3D). In both cases, fluorescent signals were detected in the nucleus and in the cell cytoplasm, with Manders’ coefficients of 0.87 and 0.82 for DENV2-mCherry and DENV2-Ct-mCherry respectively (Figure 3E). The nuclear accumulation of mCherry has been also reported for other DENV2 constructs with similar design and was most likely caused by the fusion mCherry with capsid protein or its fragment (41, 42). Previously published data indicates that the appearance of the capsid protein in the nucleus of DENV2 infected cells occurs due to the presence of nuclear localization signal that facilitates interaction with cellular proteins responsible for nuclear transport in its sequence (43, 44). Taken together, truncation of the first copy of capsid protein gene had a positive impact on the replication and genetic stability of DENV2 harboring mCherry marker without any apparent changes in subcellular distribution of the reporter. On the whole, the same approach, i.e. truncated capsid, can be used to develop flaviviruses infectious clones harboring reporters to overcome issues related to genetic instability of corresponding recombinant viruses.

### Imaging of recombinant virus-infected cells by TEM and purified viruses by Bio-AFM

To image the morphogenesis of virions of clone-derived mCherry-expressing viruses, we performed a cross analysis of resin-embedded Vero cells infected at MOI 1 by P_0_ stocks at 3 days post infection by transmission electron microscopy (TEM). Replication of flaviviruses typically causes dramatic remodeling of the endoplasmic reticulum (ER) membranes, which wrap around the viral replication factories to form viral replication organelles (VRO) (45), convoluted membranes/paracrystalline arrays (CM/PC) (46), vesicle packets (VP) used as loci for viral genome amplification, RNA translation and polyprotein processing (47, 48, 49, 50, 51, 52, 53). Coherently, infection of Vero cells with ZIKV-mCherry, DENV2-mCherry, DENV4-mCherry or KUNV-mCherry induced massive ultrastructural ER expansions and reconfigurations, cytoplasm vacuolization, formation of ER sheets and ER-derived vesicles containing electron dense viroplasm-like structures and newly formed virions (Figure 4A, left panels). CM were usually found in the center of large structures (Figure 4A), as has been previously seen in cells infected with DENV or ZIKV (18,20). VRO and VP were unambiguously seen within sub-compartments including the lumen of large cytoplasmic vacuoles, with dramatic accumulation of PC and membrane-associated virus particles, often arranged in regular arrays for DENV2 and KUNV infected cells (47, 48, 49). Tubular altered ER containing immature viral particles were also observed in close proximity to the VP (Figure 4). All these features have been commonly observed in mammalian cells producing flaviviruses (21, 23, 24).

**Figure 4.**
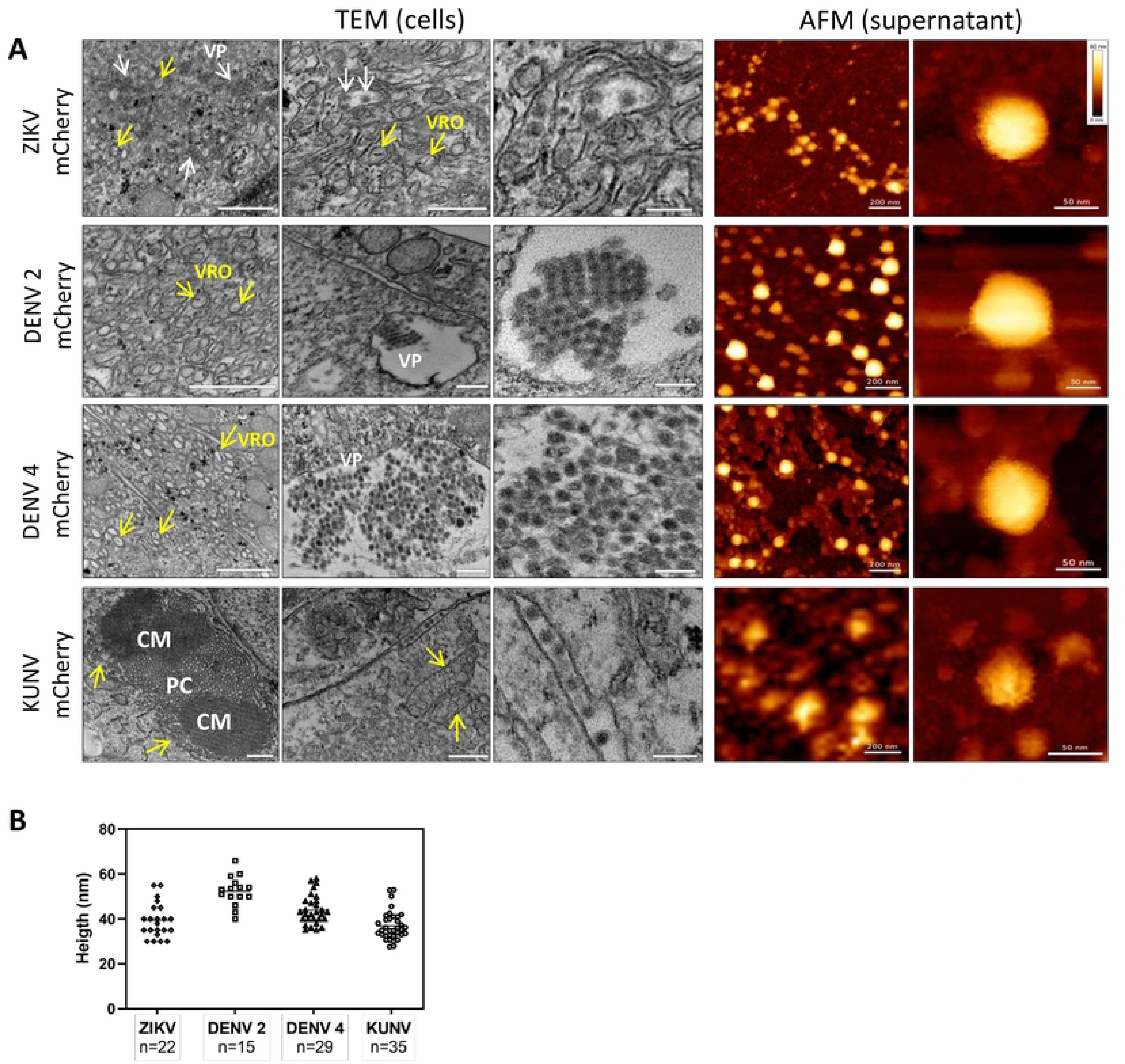
TEM and AFM images of mCherry-reporter virions. (**A**, left panel) Transmission electron microscopy of cells infected with DENV2-mCherry, DENV4-mCherry, ZIKV-mCherry and KUNV-mCherry. Replication organelles derived from the ER were visualized; CM - convoluted membranes, PC - paracrystalline arrays, VP - vesicle packets (white arrows). Localization of viral replication organelles (VRO) are shown with yellow arrows. TEM scale bar = 200nm (left and central panels) and 100 nm (right panel). (**A**, right panel) AFM images of mCherry-expressing ZIKV, DENV2, DENV4 and KUNV virions purified from cell supernatants. Virions of ZIKV-mCherry were purified by dialysis in PBS, virions of DENV2-mCherry and DENV4-mCherry were purified by Microsep advanced column from Pall Corporation in Tris-NaCl buffer and virions of KUNV-mCherry were purified by ultracentrifugation on sucrose cushion. Scale bars for AFM images are 200 nm (left panel) and 50 nm (right panel). (**B**) Height distributions of viral particles measured from cross-sections of AFM images.

Next, Bio-AFM was used to analyze the morphology and the integrity of virions released from cells infected with ZIKV-mCherry, DENV2-mCherry, DENV4-mCherry or KUNV-mCherry. Viral particles were purified from cell supernatants, resuspended/diluted in a buffer solution and smoothly adsorbed on a poly-L-lysine coated mica surface. To preserve the structural integrity of the Bio-AFM imaging was performed in a buffer, using a Bio-AFM operating in a BSL3 environment. Figure 4 (right panels) presents topographic AFM images of purified viruses with different magnifications. Interestingly, in all cases, fractions of the viral particles appeared to be clustered, reminiscent of the viral clusters observed in the ER lumen by TEM, which suggests that the virions could be released from the infected culture cells as viral “packages” or, for KUNV-mCherry, could be clustered during ultracentrifugation. Individual virions of ZIKV-mCherry, KUNV-mCherry, DENV2-mCherry and DENV4-mCherry appeared as roughly spherical particles with some angles suggesting an icosahedral arrangement, as expected from their reported cryoEM structures (53, 54, 55, 56). Viral particle height measurements obtained from cross-section analysis (Figure 4B) were also consistent with the virion sizes measured by cryoEM or previous AFM studies: 52 ± 8 nm for DENV2-mCherry (53), 44 ± 7 nm for DENV4-mCherry (56), 39 ± 9 nm for ZIKV-mCherry (54, 55) and 37 ± 8 nm for KUNV-mCherry (78). DENV4-mCherry shows morphology similar to DENV2-mCherry, with a mean diameter slightly smaller. These results suggest that modification of the viral genome (adding an extra copy of sequence encoding for capsid protein and sequence encoding mCherry, i.e. making the genome 10% larger) had no detectable impact on virion size or morphology.

### Validation of fluorescent-reporter flaviviruses for antiviral screening

To assess whether the fluorescent (or luminescent) reporter-expressing recombinant flaviviruses could be applied to high-throughput screening for antivirals in a 2D cell culture system, we chose to test the adenosine nucleoside analog NITD008, previously shown to inhibit replication of mosquito- and tick-borne flaviviruses, including WNV, DENV, YFV, ZIKV or TBEV (29, 57, 58). Cells were incubated for 2h with increasing concentrations of NITD008 and next infected with the corresponding P_0_ stocks of the reporter viruses stably expressing inserted marker protein: ZIKV-mCherry, ZIKV-NLuc, or DENV2-Ct-mCherry; ZIKV-wt or DENV2-wt infected cells were used for comparison (Figure 5). For ZIKV-mCherry and DENV-Ct-mCherry, mCherry fluorescence intensity in infected cells was quantified directly on fixed cells by fluorescence microscopy, using nuclei counting for data normalization (Figure 5A, 5E). For a sake of comparison, cells infected with wt viruses were lysed, total viral RNA were extracted and quantified by RT-qPCR (Figure 5B, 5D). For ZIKV-NLuc, bioluminescence in lysates of infected cell was quantified by conducting corresponding enzymatic assay (Figure 5C). Obtained dose-response curves allowed extracting the concentrations of NITD008 that inhibited 50% of viral infection (EC_50_). In accordance with previously reported data (59, 60), NITD008 showed, in all cases, a similar dose-dependent inhibition of virus replication, with EC_50_ values ranging from 0.75 – 1 µM for ZIKV-mCherry and ZIKV-NLuc (Figure 5B, 5C) and 0.84 µM for DENV2-Ct-mCherry (Figure 5F). Importantly, a similar EC_50_ values were obtained for wt viruses with the use of RT-qPCR quantification (Figure 5D, 5G). It allowed concluding that ZIKV-mCherry, as well as ZIKV-NLuc and DENV2-Ct-mCherry can serve as reliable tools for rapid and more direct high-throughput antiviral screening.

**Figure 5.**
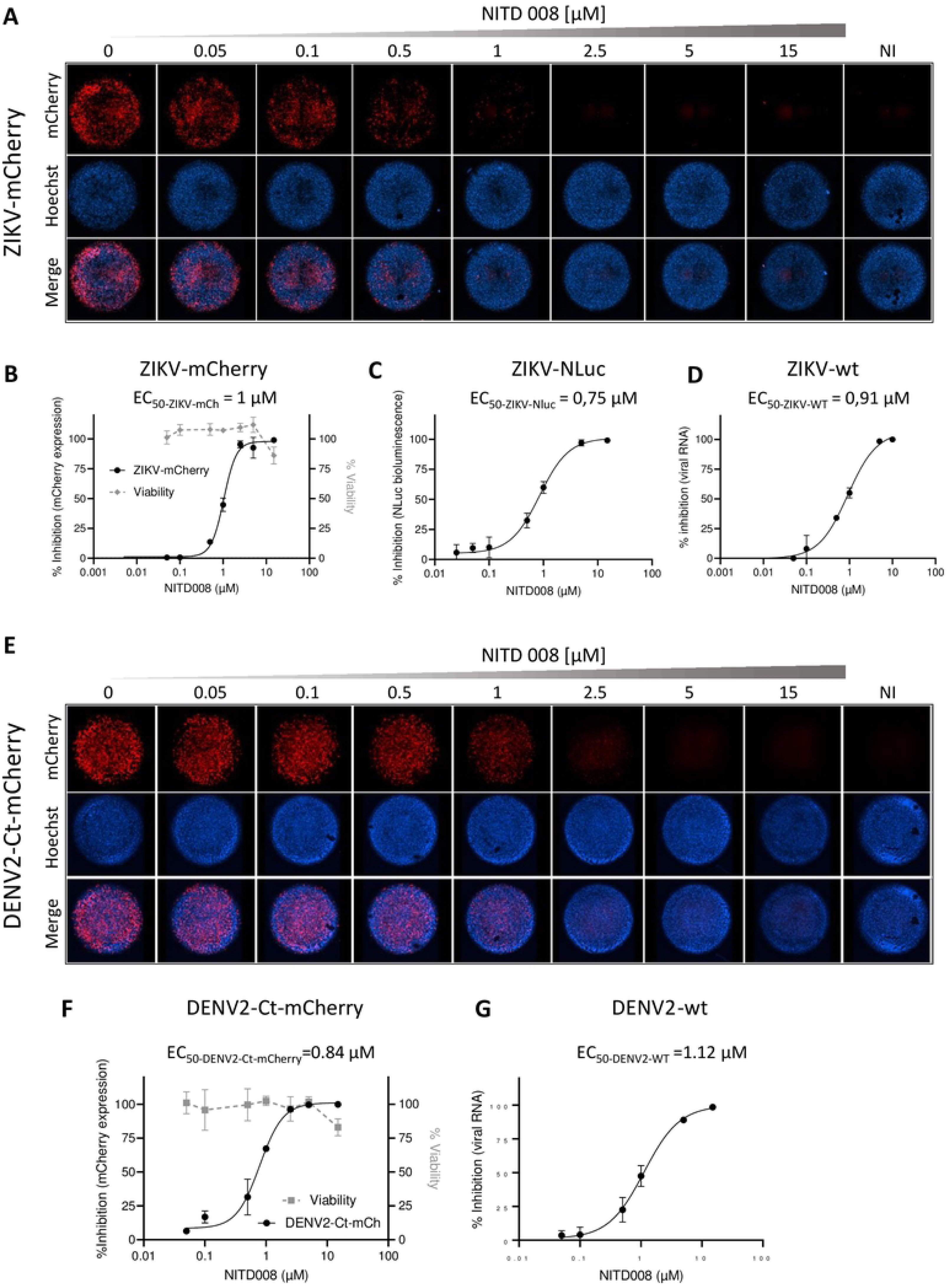
Antiviral activity of NITD008 on the reporter-expressing flaviviruses. (**A**) Dose-response of NITD008 against ZIKV-mCherry was monitored by fluorescence microscopy. Each panel corresponds to a mosaic image (6 tiles) of a single well within a 96-wells plate with Vero cells incubated with the indicated increasing concentrations (0.05, 0.1, 0.5, 1, 2.5, 5 and 15 µM) of NITD008 and infected with ZIKV-mCherry at MOI 0.05. Upper line: mCherry fluorescence, middle line: cell nuclei stained with Hoechst, bottom line: merged channels. Cell viability at 3 d.p.i. against NITD008 is shown on (**B**). Dose-response curves for ZIKV-mCherry (**B**), ZIKV-NLuc (**C**) and ZIKV-wt (**D**). (**E**) Dose-response of NIT008 against DENV2-Ct-mCh monitored by quantitative fluorescence microscopy. Vero cells were treated by increasing concentrations of NITD008 (0.05, 0.1, 0.5, 1, 2.5, 5 and 15 µM) and infected at MOI 0.1. Cell viability at 3 d.p.i. against NITD008 is shown in (**F**). Dose-response curves of NITD008 are shown for DENV2-Ct-mCherry (**F**) and DENV2-wt (**G**). The corresponding EC_50_ values are indicated in each panel.

## DISCUSSION

Over the last decades, reverse genetic approaches have allowed to advance in studying of different aspects of flavivirus replication. For this construction and manipulation of icDNA clones is essential; however, in contrast to many viruses the construction and use of these tools for flavivirus studies remains a difficult task due to the instability of the plasmids containing flavivirus cDNAs (33, 61). Despite the number of advantages of high copy number plasmids, they were generally found to be not suitable for the cloning of flavivirus icDNA since the presence of cryptic prokaryotic promoters in viral sequences leads to the synthesis of mRNAs encoding viral proteins for that are toxic for bacteria (34). An alternative approach, the use of a low copy number plasmids, often allows to overcome the instability problem of the virus cDNA by lowering the level of cryptic protein expression and consequently decreasing their cytotoxicity for *E.coli* (33). This strategy was successfully implied to obtain icDNAs for numerous flaviviruses including YFV, KUNV, ZIKV, DENV1, 2, and 4 (62, 63, 64) while DENV3 being only exception because the full-length cDNA of this virus appeared to be too toxic for maintaining in *E. coli* as a single unit (65). Our data reported above is in line with these studies as we could single plasmid icDNA clones for ZIKV, KUNV, DENV2 and DENV4 but not for DENV3. In light of this, additional approaches to resolve the stability problem have been developed. For instance, a bacterium-free methods, in particular CPER, have been successfully applied for the development of KUNV icDNA; alternatively plasmids containing flavivirus icDNAs have been stabilized by insertion of the intron into virus-derived sequences (66, 67, 68).

The use of icDNAs allows easy modification of the flavivirus genomes; however, on its own it does not permit to visualize the course and/or dynamic of the virus infection. A simple concept based on the insertion of the reporter gene(s) into the viral genome allowed to overcome such limitation. If successfully applied recombinant viruses represent valuable system for tracking and quantification of flavivirus replication *in vitro* and, potentially, also *in vivo*. The findings obtained in the current study confirmed that such viral constructs can be used for high-content antiviral screening as well as for acceleration of antiviral molecules discovery. However, generation of the reporter viruses containing oxGFP, mCherry, or NLuc markers does encounter another problem – genetic instability of rescued recombinant viruses seen as loss of the reporters due to the recombination events during virus replication and selection for faster replicating recombinants (lacking reporter gene) during virus propagation and passaging.

Here we observed that recombinant flaviviruses harboring NLuc marker were somewhat typically more stable than the ones carrying mCherry or oxGFP markers (Figure 2). Presumably, this was caused by smaller size of NLuc insertion (sequence encoding NLuc has length 513 bp while that for mCherry is 708 bp long). This is consistent with our previous studies with ZIKV where presence of additional 228 bp insertion encoding ubiquitin resulted in marked destabilization of viruses encoding for mCherry or GFP reporters (29). Such effect was not observed for ZIKV encoding NLuc suggesting existence of rather well-defined upper limit for size of insert that is tolerated by ZIKV. Data presented in this study suggest that this limit is different for each flavivirus and likely even for sub-type of virus as seen on example of DENV2 and DENV4. Interestingly, the stability assay revealed that fluorescent markers were better tolerated by ZIKV than other tested flaviviruses (Figure 2). We observed that previously reported data about stability of marker in ZIKV-RLuc (64), DENV2-mCherry (26) and DENV2-GFP (70) is coherent to our data. Thus, the findings obtained in the current study clearly underlined the high level of instability of all of DENV2 and KUNV variants harbouring reporters (Figure 2).

It can be speculated that depending on the viral species lengthening of the genome due to the insertions may have a different negative impact on cyclization of the genome and/or packaging of the viral RNA into the capsid without having a detrimental effect on the virus virulence. Noteworthily, one of the recent discoveries demonstrated that JEV had the ability to pack the genomic RNA of much larger in length (up to 15 kb) but the increase of genome length was accompanied by a decreased efficiency of RNA replication (69). Our data indicates that an absolute speed of the replication is unlikely the cause, since KUNV variants replicate faster than these of ZIKV or DENV4 (Figure 1). In the recent study, comparison of stability of NLuc insertion for 10 passages was performed for DENV1-4, ZIKV, YFV, JEV. This study revealed a clear correlation between the marker stability and length of the 5’ copy of capsid-encoding sequence (36). Our observation that a reduction of the length of 5’ copy of capsid-encoding sequence in the DENV2-mCherry has clear positive impact on the stability of the marker expression over passages (Figure 3) is coherent with this data.

Earlier reports also confirmed that some viruses may tolerate less than 10% of an increase in their genome size during packaging into viral capsids highlighting the importance of the relationship between the size of the viral genome and capsid formation. In this regard it should be mentioned that we could not find any correlation between the size of the RNA genome and recombinant reporter virus stability as sizes of ZIKV (10807 nt), KUNV (11022 nt), DENV2 (10723 nt), and DENV4 (10649 nt) genomes are similar. It may be the ratio of the size of genome to the size of the capsid or the virion that is important - not all viruses may have the same capsid size and the compaction of the viral genome would be higher in a smaller capsid. Clearly, more detailed studies are needed to discover precisely the mechanism(s) responsible for disadvantage (and therefore counterselection) of flaviviruses with increased length of genomic RNA.

Capsid protein is known to be a central element in flavivirus assembly process which due to the precise physical interactions with the viral genome stabilize the capsid (71). The anchor domain (α5 helix) of the capsid possess a key function in the virion assembly and in case of its absence, the capsid dimers may remain “locked” within the RNA core impeding the correct formation of the virions. What is more interesting, controlled and timeliness manner of the capsid α5 helix processing was shown as one of the crucial factors necessary for incorporation of the nucleocapsid into the virions. Subsequently, α5 helix is removed from the capsid protein but recent investigations demonstrated the opposite for ZIKV - α5 helix is retained for some of capsid subunits present in virions indicating other possible (additional) functions in the assembly process (71, 72). Additionally, the threshold of the positive charges localized in the N-terminal part of capsid protein which are necessary for the correct formation of the virions may be different and thus may affect the stability of flaviviruses to a different extend. Taken it together, requirements for the involvement of capsid protein in the coordination of the virion formation are different which in turn may influence the overall stability of the increased length of the genome or its packaging. Moreover, the functional properties of the RNA elements present within the first 38 codons in the 5’ copy of capsid gene were found to be different among flaviviruses (39). It indicates that shortening of this sequence may not affect viral fitness to the same extent for all flaviviruses.

Flavivirus particles sizes determined so far by cryoEM are for ZIKV ∼49 nm (73), DENV2 ∼50 nm (74, 75), DENV4 ∼ 48 nm (56) and KUNV ∼50 nm (76). The AFM measurements performed in the course of the study (Figure 4) with reporter viruses containing mCherry insertion in their genomes are consistent for DENV2 (∼ 52 nm) and DENV4 (∼ 44 nm) particle sizes; interestingly somewhat smaller particles’ sizes were measured for ZIKV (∼ 40 nm) and KUNV (37 nm). This suggests a slight deformation of ZIKV and KUNV particles during the AFM experiment that could be due either to their adsorption on the surface or to the force applied by the AFM tip (or both). On the other hand, DENV2 and DENV4 are maybe more structurally stable in these conditions offering a comparable particle sizes between cryoEM and AFM imaging. However, our results suggest an absence of correlation between particle size and marker stability, as exemplified by ZIKV. Therefore, the size of the virion may be affected – to some extend - by the envelope organization rather than the capsid/genomic RNA size. This leaves the question open. In addition, packages of secreted virions (Figure 4) have not been described so far, to our knowledge, and further explorations are required to attribute this phenomenon to particles preparation for imaging or to a biological behavior.

How does the loss of marker occur? Here we did not analyze the mechanism directly but some conclusions can be based on observed speed of the marker loss. Previous studies indicate that initial loss of marker occurs by in-frame deletions in the reporter encoding sequence. In most cases loss of reporter did not occur (or, at least did not become detectable) during rescue; instead it occurred, often very rapidly, but during passaging of rescued virus. It is thus very likely that the main mechanism how markerless viruses become dominant is not the loss of the marker itself (the timing or frequency of such an event) during viral rescue and replication. Instead, our data is more coherent with hypothesis that difference between growth kinetics of wt virus and virus with reporter may be the key: the smaller the growth advantage of wt virus the most stable are its variants with reporter. The significant growth advantage of wt viruses most likely extends also to viruses harboring deletion in marker region and allows rapid outcompeting the ones that have maintained marker.

Given these considerations, the new approaches directed to increasing of the stability of recombinant flavivirus genomes preserving the properties and virulence similar to the wild type virus are needed. Up to date, several strategies have been developed and tested. One of them is utilization of recombination-dependent lethal mutations. Introduction of such a change into the capsid region of ZIKV and YFV is capable to increase the stability and prevent the formation of defective flavivirus virion in case recombination events. At the same time, the use of split reporter proteins can also be considered to be one of the most promising approach due to it already proven robustness and effectiveness of these reporters. Finally, recent study showed that a point nucleotide mutation (T142C) in the 5’ CS of JEV, WNV and DENV leads to the mutation of the capsid protein at the position 16 (M16T) which in turn assist to genome cyclization with the following stabilization of the reporter-harboring viruses (77). The question that remains open is whether the combination of several previously mentioned approaches can increase the stabilizing effect.

## ACKNOWLEDGMENTS

We thank Yannick Simonin from University of Montpellier (France) for helpful discussions, Benoît Bordignon from MRI (CNRS Montpellier, France) and Julien Kissenberger from ZEISS France for help with data acquisition and image analysis. The BSL3 Bio-AFM was funded by the REDSAIM program by Montpellier University, the BSL3 Cell-Discoverer 7 microscope was funded by Occitanie FEDER. We thank the CNRS and Montpellier University for funding. AM and LC were supported by project grant [PRG1154] from Estonian Research Council.

## AUTHOR CONTRIBUTIONS

LC, AT, NG, MH, MV, AN, CCB, AM and SL performed experiments. LC and AM performed construction of icDNA clones; LC, AT, NG, MH, CCB performed virus production and purification, viral titers, immunoblots, infection kinetics, FFA and data analysis; AT and SL performed AFM analysis; MV and SL performed fluorescence microscopy imaging and analysis; AN performed TEM imaging; NG, SL, AM and DM conceived, directed, and supervised the study. SL, LC, NG, AT, AM and DM wrote and edited the manuscript. AM and DM raised funding.

## MATERIAL AND METHODS

### Cell culture

Vero cells (African green monkey kidney cells, ATCC CCL-81) were grown in Dulbecco’s modified Eagle’s medium (DMEM, Lonza) supplemented with 10% foetal bovine serum (FBS, Gibco), 100 U/mL of penicillin and 100 µg/mL streptomycin at 37°C with 5% CO_2_.

### Design and assembly of wt and reporter harboring icDNA clones of flaviviruses

Construction of icDNA clone of ZIKV (Brazilian isolate), designated as ZIKV-wt, and its variants harboring mCherry and NLuc reporters (ZIKV-mCherry and ZIKV-NLuc) has been previously described (32). To obtain ZIKV expressing green fluorescent marker, the NLuc was replaced with oxidation resistant GFP marker (58), obtained clone was designated ZIKV-oxGFP. The same cloning strategy was applied for construction of icDNA clones of DENV2, DENV4 and KUNV. Briefly, synthetic DNA fragments were obtained from Twist Bioscience (San Francisco, CA, USA) and Genscript (New Jersey, USA). The assembly strategy included five steps in which the synthetic fragments were consequentially cloned into the single copy pCCI-Bac plasmid with SP6 promoter placed upstream of region corresponding to the 5’ end of the virus genome. The oxGFP, mCherry, and NLuc marker genes were cloned between two copies of capsid sequences in the structural region of the modified genome as described before (32). The obtained clones were designated as DENV2-wt (-oxGFP, -mCherry, -NLuc), DENV4-wt (-oxGFP, -mCherry, -NLuc), and KUNV-wt (-oxGFP, -mCherry, -NLuc). To reduce the loss of mCherry marker, a 228 bp deletion, removing codons 39-114 of the first (native) copy of the region encoding for capsid protein of DENV2 was introduced into DENV2-mCherry using PCR-based mutagenesis and subcloning procedures; resulting clone was designated as DENV2-Ct-mCherry. Cloning procedures and amplification of the obtained plasmids containing the icDNAs of the viruses was done in *E. coli* EPI300 cells (LGC Biosearch Technologies, UK). Sequences of all obtained plasmids were confirmed using Sanger sequencing and are available form authors upon request.

### *In vitro* transcription and virus rescue

10 µg of icDNA plasmids of DENV2-wt, DENV4-wt, ZIKV-wt, KUNV-wt and their reporter-containing variants were linearized using AgeI-HF enzyme (NEB, United States) prior *in vitro* transcription. The linearized DNAs were purified using Monarch DNA cleanup kit (NEB, United States) and the capped RNA transcripts were synthesized using the SP6 mMessage mMachine kit (Invitrogen, United States) following manufacturer’s instructions.

All virus studies were conducted under biosafety level 3 facility in CNRS CEMIPAI Montpellier. Vero cells were transfected with obtained RNA transcripts using Lipofectamine 2000 (Invitrogen) reagent. Briefly, transfection mixtures were incubated at room temperature for 5 min and added to Vero cells monolayer with following incubation for 5-6 h at 37°C. The cells were then washed in 1x PBS (Eurobio, France) and incubated in growth media at 37°C. Supernatants (P_0_ stocks) were collected at 7 to 15 days post transfection, clarified by centrifugation at 1000 x g for 10 min, aliquoted and stored at −80°C.

### Focus Forming Assay (FFA)

Viral supernatants were titrated by the end-point titration method. Vero cells were seeded on 96-well plates at 10,000 cells per well and incubated for 4h at 37°C. Cells were infected with virus dilutions (from 10^-1^ to 10^-9^) prepared in DMEM supplemented with 2% FBS and 1%Penicillin/Streptomycin mixture (Lonza Biosciences). Infected cells were incubated at 37°C for 7 to 15 days. The medium was then removed, cells were washed once with PBS and fixed with 4% paraformaldehyde (PFA, Fisher) for 30 min at room temperature. After removal of PFA, cells were washed with PBS and kept at 4°C until staining. For the staining, cells were permeabilized with PBS containing 0.1% Triton X-100 (Sigma Aldrich, France) for 5 min and blocked with PBS containing 2% FBS and 0.05% Tween20 (Sigma Aldrich, France) for 1h. Cells were washed once with PBS and incubated with a mouse pan-flavivirus anti-Env antibody (Mab 4G2, Novus Biologicals NBP2) for 1-2 h at room temperature (1:1000) for viral titration or an anti-GFP antibody (A11122, Invitrogen) for measurement of oxGFP-positive cells. Following this, cells were washed 3 x 10 min with PBS containing 0.1% Tween20 and then incubated in the dark with a fluorescent goat anti-mouse secondary antibody conjugated with DyLight 800 (Invitrogen, SA5-35521) for 1-2h at room temperature. Finally, cells were washed 3 x 10 min with PBS containing 0.1% Tween20 and the fluorescence was recorded with a microplate reader (Odyssey, Li-Cor Biosciences, United States). Viral titers were calculated using the Spearman & Kärber algorithm.

### Genetic stability assay

Vero cells were seeded on 6-well plates at 300,000 cells per well. Cells were infected by adding 500 µL of P_0_ supernatants corresponding to each of the rescued marker-expressing viruses. After incubation for 2h, the viral supernatants were removed, cells were washed once with PBS and 2mL of fresh growth medium was added. Cells were incubated for 5-15 days at 37°C. Viral supernatants (P_1_ stocks) were collected and used for following passages performed as described above; supernatants from each passage (P_2_, P_3_ and P_4_) were collected. FFA was used to quantify the titers of the rescued viruses in collected virus stocks. mCherry expression was determined based on the signal intensity measurement using Cellomics ArrayScan VTI microscope. oxGFP expression was detected by immunofluorescence assay using an anti-GFP antibody (A11122, Invitrogen). For the measurement of NLuc activity, a second 96-well plate was infected with P1 to P4 stocks according to the end-point titration method and NLuc was measured with the Nano-Glo Luciferase Assay System (Promega) after cell lysis with 1X passive lysis buffer (E1941, Promega), on the Envision (Perkin Elmer plate reader). The percentage of the virus-infected cells expressing reporters was calculated based on the ratio of the number of wells positive for the marker (mCherry, NLuc or oxGFP) and the number of wells positive for the virus (detected by staining with 4G2 antibody).

### Western blot

8 mL of viral supernatant on a 3 mL of 20% sucrose cushion were ultracentrifugated for 3 h at 100,000 x g at 4°C using an Optima L80-XP instrument (Beckman Coulter). After ultracentrifugation, the liquid was discarded, the tube was dried with a Kimtech paper and the pellet was resuspended in 80 µL of TNE buffer (10 mM TrisHCl (pH 7.0), 100mM NaCl, 1mM EDTA). Obtained samples were denatured in a 1xLaemmli buffer at 95°C for 10 min and proteins separated using electrophoresis on a 4-15% mini-protean-TGX-precast-gel (BioRad). Proteins were then transferred on nitrocellulose membranes by a semi-dry method using transblot turbo transfer system (BioRad). The membranes were blocked with 5% skimmed milk powder in PBS-Tween20 for 30 min and incubated overnight at 4°C with either virus-specific primary capsid antibody (rabbit anti-capsid ZIKV GTX134186 in dilution 1:10000; mouse anti-capsid DENV GTX633632 in dilution 1:3,000; rabbit anti-capsid WNV GTX131947-S in dilution 1:100; all antibodies from GeneTex) or primary anti-pan flavivirus envelope antibody (mouse anti D1-4G2-4-15 (4G2) by NovusBio NBP2-52666 in dilution 1:1000). Membranes were rinsed 3 x 10 min in PBS-Tween20 prior to incubation with appropriate secondary antibody (anti-mouse DyLight 800 and anti-rabbit DyLight 800 conjugated antibodies SA5-35521 or SA5-35571 from Invitrogen) for 2 h at room temperature. Membranes were rinsed 3 x 10 min in PBS-Tween20 before reading on a Li-Cor Odyssey scanner 9120.

### Immunofluorescence analysis

200,000 Vero cells grown on 35 mm dishes (Fluorodish, Fisher) and infected at multiplicity of infection (MOI) 0.05 with P_0_ (collected at day 5 post transfection) stocks of all analyzed viruses. Infected cells were incubated in for three (ZIKV, DENV2, KUNV) or four (DENV4) days after which cells were fixed with 4% PFA (Fisher) for 30 min at 20°C. Fixed cells were rinsed with PBS, permeabilized with 0.1% Triton X-100 diluted in PBS for 5 min and blocked with 2% BSA. After, the cells were incubated in PBS supplemented with the pan-flavivirus anti-Env monoclonal antibody 4G2 (NovusBio, dilution 1:1000) at the presence of 0.05% Saponin (Sigma-Aldrich, France) for 1h, washed 3 times with PBS and incubated with anti-mouse secondary antibodies conjugated with Alexa647 (ab150107, Abcam, 1:1000 dilution) or Alexa488 (A21202, Invitrogen, 1:10000 dilution) 2 h at 4°C in the dark. Nuclei were stained with 10µg/mL Hoechst 33342 (Invitrogen) for 15 min. Stained cells were washed with PBS and epifluorescence and confocal images were acquired using a Cell-Discoverer 7 microscope (Carl Zeiss SAS, France) at 10x and 25× magnification. Mander’s coefficients were calculated based on the images obtained using Zen Blue software (Zeiss).

### RT-qPCR

The cells were infected with ZIKV and DENV2 and lysed using the Luna Cell Ready Lysis reagent (New England Biolabs, UK). Quantification of viral RNA was performed from cell lysates using the virus specific primers and their levels were normalized to mRNA of GAPDH (Table 1). For these analyses Luna Universal One-Step RT-qPCR Kit (New England Biolabs) on a CFX opus real-time 384 system (BioRad) were used. Cycling conditions were as follows: reverse transcription at 55°C for 15 minutes, followed by initial polymerase activation at 95°C for 1 min, and then 45 cycles of denaturation at 95°C for 10 seconds and annealing/extension at 60°C for 45 seconds. The calibration of the assay was performed with control plasmids containing sequences encoding ZIKV NS1 and DENV2 capsid proteins (Eurofins Genomics, Germany; GenBank accession number KU365778.1 and U87411.1 respectively).

**Table 1:** Primers used for RT-qPCR for viral RNA quantification.

### Antiviral assay

NITD008 (Bio-Techne 6045/1) was initially dissolved in 100% dimethyl sulfoxide (DMSO D8418, SIGMA) at 10 mM and subsequently diluted in DMEM to the desired concentrations (0.05 to 15 µM). 0.5% DMSO was set as vehicle control. Cells were incubated with increasing concentrations of NITD008 for 2 h prior infection. For cell fluorescence counting, 9,000 cells per well were cultured in black opaque 96-well Microplate (PerkinElmer), infected with virus stock at an MOI 0.01 (ZIKV-mCherry) or MOI 0.05 (DENV2-Ct-mCherry) and prepared for imaging at 72 h post infection. Briefly, cells were fixed with 4% PFA and stained with 10 µg/mL Hoechst 33342 for 15 min and washed with PBS. 3×2 image tiles acquisitions per well were performed at 2.5 magnification in wide-field on a Cell-Discoverer 7 microscope (Carl Zeiss SAS, France). Image analysis was performed using ZEN Blue software for segmenting of the stained nuclei and mCherry positive cells. The number of nuclei were sorted per well. The sum of fluorescence intensity was weighted against the normalized number of nuclei. For measurement of NLuc activity 20,000 cells were plated on transparent 96-well plate and infected with ZIKV-NLuc at MOI 0.05. Luminescence was measured at 1 day post infection using the Nano-Glo Luciferase Assay System (Promega) on the Envision plate reader after cell lysis with 1X passive lysis buffer (E1941, Promega). Results were normalized to the total protein amount per well determined using a BCA protein assay kit (Pierce). For wt viruses, 20,000 cells were cultured in transparent 96-well plate and infected with DENV2-wt or ZIKA-wt at an MOI of 0.01 and 0.05 respectively. 72 h post-infection, cells were lysed and total viral RNA extracted and quantified by RT-qPCR as described above. In all cases, the 50% effective concentration (EC_50_) was calculated by fitting the dose-response curves traced from 4-parameter nonlinear regression with GraphPad Prism v9.0, based on the following calculations:

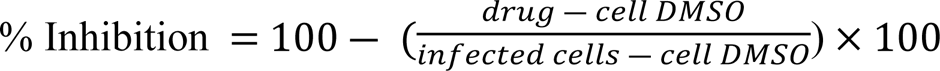

### Transmission Electron Microscopy (TEM)

Vero cells were infected at an MOI 1, incubated for 3 days and fixed with 2.5% glutaraldehyde in PHEM buffer (PIPES 60mM, HEPES 25mM, EGTA 10mM, MgCl2 2mM, all provided by MERCK-SIGMA, Germany), post fixed in 1% OsO_4_ / 0.8% K_4_Fe (CN)_6_ then dehydrated in successive ethanol bathes (50/70/90/100%). Samples were then infiltrated in propylene oxide (MERCK-SIGMA, Germany)/EMbed812 (EMS, USA) mixes, embedded in EMbed812 and polymerized at 60°C. Ultrathin sections (70nm) were cut using a PowerTome XL ultramicrotome (RMC, Tucson, AZ, USA), stained in 0.2% OTE/lead citrate and observed on a Tecnai G2 F20 (200kV FEG) TEM at the platform Plateau de Microscopie Electronique COMET, INM.

### Atomic Force Microscopy (AFM)

AFM imaging was performed on a JPK-Bruker Nanowizard IV XP atomic force microscope (JPK BioAFM, Bruker Nano GmbH, Berlin, Germany) operating in BSL-3 (37). Flavivirus virions were imaged in imaging buffer (10mM Tris-HCl pH 7.5, 100mM NaCl) shortly after being adsorbed on freshly cleaved muscovite surface (mica grade v1, Ted Pella) glued on a glass slide. Before virus adsorption, the mica was functionalized with 100 µL Poly-L-Lysine 0.1% (Sigma P8920) for 10 min, then washed three times in imaging buffer. A 3D printed plastic O-ring was glued around the mica to form a small liquid cell. 20 µL of viral samples were deposited on the functionalized mica from 15 min to 1 h, and the volume was completed to 200 µL with the imaging buffer. AFM topographic images were obtained using the quantitative imaging (QI) mode using qp-Bio-AC CB2 (Nanosensors), BL-AC40TS (Olympus) or MLCT-Bio (Bruker) cantilevers. Before each set of acquisitions, the sensitivity and spring constant of the cantilever were calibrated (thermal noise method). The applied force was kept at 150 to 200 pN, 100 nm Z-length and 20 msec/pixel speed. Using the JPK SPM-data processing software, images were flattened with a polynomial/histogram line fit. Low-pass Gaussian and/or median filtering was applied to remove minor noise from the images. The Z-color scale in all image is given as relative after processing. Particle height analysis, based on the height (measured) channel of the QI mode, was performed using the cross section of the analysis software to calculate the maximal central height on each particle.

### Statistical analysis

Statistical analysis was performed with GraphPad Prism 9.3.0 software. The results of marker expression were represented with error bars indicating the standard deviation (SD).

## Supplementary Figures

**Supplementary Figure S1: Detection of envelope and capsid proteins in purified recombinant flavivirus particles.** (**A**) Viral supernatants were collected, virus particles were ultracentrifugated and lysed in SDS loading butter. Obtained samples were separated by SDS-PAGE and viral proteins were detected using immunoblotting with either pan-*Flavivirus* envelope (4G2) or specific *Flavivirus* anti-capsid antibodies. NI – not infected; * designates nonspecific signal or envelope multimers in sample obtained from DENV4 particles. (**B**) Viral titers measured by FFA (in focus forming units (FFU)/mL) in the viral supernatants used to prepare the samples analyzed in (**A**).

**Supplementary Figure S2**: **Tracking the intensity of the mCherry fluorescent signal over time** for ZIKV-, KUNV-, DENV2- and DENV4-mCherry expressing constructs in transfected Vero cells from day 1 up to day 12. Scale bars: 100 nm.

## REFERENCES

1. Transmission cycles, host range, evolution and emergence of arboviral disease. Nat Rev Microbiol. 2004; 2(10): 789–801. doi:10.1038/nrmicro1006

2. Pierson TC, Diamond MS. The continued threat of emerging flaviviruses. Nat Microbiol. 2020 Jun;5(6):796–812. doi: 10.1038/s41564-020-0714-0. Epub 2020 May 4. PMID: 32367055; PMCID: PMC7696730.

3. Choi, K.H. The Role of the Stem-Loop A RNA Promoter in Flavivirus Replication. Viruses 2021, 13, 1107. https://doi.org/10.3390/v13061107

4. Mazeaud C, Freppel W, Chatel-Chaix L. The Multiples Fates of the Flavivirus RNA Genome During Pathogenesis. Front Genet. 2018 Dec 4;9:595. doi: 10.3389/fgene.2018.00595. PMID: 30564270; PMCID: PMC6288177.

5. Pan Y, Cai W, Cheng A, Wang M, Yin Z, Jia R. Flaviviruses: Innate Immunity, Inflammasome Activation, Inflammatory Cell Death, and Cytokines. Front Immunol. 2022 Jan 28;13:829433. doi: 10.3389/fimmu.2022.829433.

6. Guabiraba R, Ryffel B. Dengue virus infection: current concepts in immune mechanisms and lessons from murine models. Immunology. 2014 Feb;141(2):143–56. doi: 10.1111/imm.12188.

7. Gould E.A., Solomon T. Pathogenic flaviviruses. Lancet. 2008;371:500–509.

8. Khanam A, Gutiérrez-Barbosa H, Lyke KE, Chua JV. Immune-Mediated Pathogenesis in Dengue Virus Infection. Viruses. 2022 Nov 21;14(11):2575. doi: 10.3390/v14112575.

9. Barbi L, Coelho AVC, Alencar LCA, Crovella S. Prevalence of Guillain-Barré syndrome among Zika virus infected cases: a systematic review and meta-analysis. Braz J Infect Dis. 2018 Mar-Apr;22(2):137-141. doi: 10.1016/j.bjid.2018.02.005.

10. Lima MES, Bachur TPR, Aragão GF. Guillain-Barre syndrome and its correlation with dengue, Zika and chikungunya viruses infection based on a literature review of reported cases in Brazil. Acta Trop. 2019 Sep;197:105064. doi: 10.1016/j.actatropica.2019.105064.

11. Wen Z, Song H, Ming GL. How does Zika virus cause microcephaly? Genes Dev. 2017 May 1;31(9):849–861. doi: 10.1101/gad.298216.117.

12. Colpitts TM, Conway MJ, Montgomery RR, Fikrig E. West Nile Virus: biology, transmission, and human infection. Clin Microbiol Rev. 2012 Oct;25(4):635–48. doi: 10.1128/CMR.00045-12.

13. van Leur SW, Heunis T, Munnur D, Sanyal S. Pathogenesis and virulence of flavivirus infections. Virulence. 2021 Dec;12(1):2814–2838. doi: 10.1080/21505594.2021.1996059.

14. Che, P., Wang, L. & Li, Q. The development, optimization and validation of an assay for high throughput antiviral drug screening against Dengue virus. International journal of clinical and experimental medicine 2, 363 (2009).

15. Xu HT, Colby-Germinario SP, Hassounah SA, Fogarty C, Osman N, Palanisamy N, Han Y, Oliveira M, Quan Y, Wainberg MA. Evaluation of Sofosbuvir (β-D-2’-deoxy-2’-α-fluoro-2’-β-C-methyluridine) as an inhibitor of Dengue virus replication. Sci Rep. 2017 Jul 24;7(1):6345. doi: 10.1038/s41598-017-06612-2.

16. McCormick KD, Liu S, Jacobs JL, Marques ET Jr, Sluis-Cremer N, Wang T. Development of a robust cytopathic effect-based high-throughput screening assay to identify novel inhibitors of dengue virus. Antimicrob Agents Chemother. 2012 Jun;56(6):3399–401. doi: 10.1128/AAC.06425-11.

17. Gurukumar KR, Priyadarshini D, Patil JA, Bhagat A, Singh A, Shah PS, Cecilia D. Development of real time PCR for detection and quantitation of Dengue Viruses. Virol J. 2009 Jan 23;6:10. doi: 10.1186/1743-422X-6-10.

18. Faye O, Faye O, Diallo D, Diallo M, Weidmann M, Sall AA. Quantitative real-time PCR detection of Zika virus and evaluation with field-caught mosquitoes. Virol J. 2013 Oct 22;10:311. doi: 10.1186/1743-422X-10-311.

19. Wilson HL, Tran T, Druce J, Dupont-Rouzeyrol M, Catton M. Neutralization Assay for Zika and Dengue Viruses by Use of Real-Time-PCR-Based Endpoint Assessment. J Clin Microbiol. 2017 Oct;55(10):3104–3112. doi: 10.1128/JCM.00673-17.

20. Cruz DJ, Koishi AC, Taniguchi JB, Li X, Milan Bonotto R, No JH, Kim KH, Baek S, Kim HY, Windisch MP, Pamplona Mosimann AL, de Borba L, Liuzzi M, Hansen MA, Duarte dos Santos CN, Freitas-Junior LH. High content screening of a kinase-focused library reveals compounds broadly-active against dengue viruses. PLoS Negl Trop Dis. 2013;7(2):e2073. doi: 10.1371/journal.pntd.0002073.

21. Shum D, Smith JL, Hirsch AJ, Bhinder B, Radu C, Stein DA, Nelson JA, Früh K, Djaballah H. High-content assay to identify inhibitors of dengue virus infection. Assay Drug Dev Technol. 2010 Oct;8(5):553–70. doi: 10.1089/adt.2010.0321.

22. Koishi AC, Suzukawa AA, Zanluca C, Camacho DE, Comach G, Duarte Dos Santos CN. Development and evaluation of a novel high-throughput image-based fluorescent neutralization test for detection of Zika virus infection. PLoS Negl Trop Dis. 2018 Mar 15;12(3):e0006342. doi: 10.1371/journal.pntd.0006342.

23. Schoggins JW, Dorner M, Feulner M, Imanaka N, Murphy MY, Ploss A, Rice CM. Dengue reporter viruses reveal viral dynamics in interferon receptor-deficient mice and sensitivity to interferon effectors in vitro. Proc Natl Acad Sci U S A. 2012 Sep 4;109(36):14610–5. doi: 10.1073/pnas.1212379109.

24. Zou G, Xu HY, Qing M, Wang QY, Shi PY. Development and characterization of a stable luciferase dengue virus for high-throughput screening. Antiviral Res. 2011 Jul;91(1):11–9. doi: 10.1016/j.antiviral.2011.05.001.

25. Zhang JW, Wang H, Liu J, Ma L, Hua RH, Bu ZG. Generation of A Stable GFP-reporter Zika Virus System for High-throughput Screening of Zika Virus Inhibitors. Virol Sin. 2021 Jun;36(3):476–489. doi: 10.1007/s12250-020-00316-0.

26. Li LH, Kaptein SJF, Schmid MA, Zmurko J, Leyssen P, Neyts J, Dallmeier K. A dengue type 2 reporter virus assay amenable to high-throughput screening. Antiviral Res. 2020 Nov;183:104929. doi: 10.1016/j.antiviral.2020.104929.

27. Fischl W, Bartenschlager R. High-throughput screening using dengue virus reporter genomes. Methods Mol Biol. 2013;1030:205–19. doi: 10.1007/978-1-62703-484-5_17.

28. Netsawang J, Noisakran S, Puttikhunt C, Kasinrerk W, Wongwiwat W, Malasit P, Yenchitsomanus PT, Limjindaporn T. Nuclear localization of dengue virus capsid protein is required for DAXX interaction and apoptosis. Virus Res. 2010 Feb;147(2):275–83. doi: 10.1016/j.virusres.2009.11.012.

29. Mutso M, Saul S, Rausalu K, Susova O, Žusinaite E, Mahalingam S, Merits A. Reverse genetic system, genetically stable reporter viruses and packaged subgenomic replicon based on a Brazilian Zika virus isolate. J Gen Virol. 2017 Nov;98(11):2712–2724. doi: 10.1099/jgv.0.000938.

30. Lyonnais, et al., 2021

31. Ruggli N, Rice CM. Functional cDNA clones of the Flaviviridae: strategies and applications. Adv Virus Res. 1999;53:183–207. doi: 10.1016/s0065-3527(08)60348-6.

32. Zheng, X., Tong, W., Liu, F. et al. Genetic instability of Japanese encephalitis virus cDNA clones propagated in Escherichia coli. Virus Genes 52, 195–203 (2016). https://doi.org/10.1007/s11262-016-1289-y

33. Aubry F, Nougairède A, Gould EA, de Lamballerie X. Flavivirus reverse genetic systems, construction techniques and applications: a historical perspective. Antiviral Res. 2015 Feb;114:67–85. doi: 10.1016/j.antiviral.2014.12.007.

34. Pu SY, Wu RH, Yang CC, Jao TM, Tsai MH, Wang JC, Lin HM, Chao YS, Yueh A. Successful propagation of flavivirus infectious cDNAs by a novel method to reduce the cryptic bacterial promoter activity of virus genomes. J Virol. 2011 Mar;85(6):2927–41. doi: 10.1128/JVI.01986-10.

35. Baker C, Xie X, Zou J, Muruato A, Fink K, Shi PY. Using recombination-dependent lethal mutations to stabilize reporter flaviviruses for rapid serodiagnosis and drug discovery. EBioMedicine. 2020 Jul;57:102838. doi: 10.1016/j.ebiom.2020.102838.

36. Baker C, Shi PY. Construction of Stable Reporter Flaviviruses and Their Applications. Viruses. 2020 Sep 25;12(10):1082. doi: 10.3390/v12101082.

37. Mutso M, Saul S, Rausalu K, Susova O, Žusinaite E, Mahalingam S, Merits A. Reverse genetic system, genetically stable reporter viruses and packaged subgenomic replicon based on a Brazilian Zika virus isolate. J Gen Virol. 2017 Nov;98(11):2712–2724. doi: 10.1099/jgv.0.000938.

38. Baker C, Liu Y, Zou J, Muruato A, Xie X, Shi PY. Identifying optimal capsid duplication length for the stability of reporter flaviviruses. Emerg Microbes Infect. 2020 Dec;9(1):2256–2265. doi: 10.1080/22221751.2020.1829994.

39. Volkova E, Tsetsarkin KA, Sippert E, Assis F, Liu G, Rios M, Pletnev AG. Novel Approach for Insertion of Heterologous Sequences into Full-Length ZIKV Genome Results in Superior Level of Gene Expression and Insert Stability. Viruses. 2020 Jan 3;12(1):61. doi: 10.3390/v12010061.

40. Samsa MM, Mondotte JA, Caramelo JJ, Gamarnik AV. Uncoupling cis-Acting RNA elements from coding sequences revealed a requirement of the N-terminal region of dengue virus capsid protein in virus particle formation. J Virol. 2012 Jan;86(2):1046–58. doi: 10.1128/JVI.05431-11.

41. Bulich R, Aaskov JG. Nuclear localization of dengue 2 virus core protein detected with monoclonal antibodies. J Gen Virol. 1992 Nov;73 (Pt 11):2999–3003. doi: 10.1099/0022-1317-73-11-2999.

42. Welsch S, Miller S, Romero-Brey I, Merz A, Bleck CK, Walther P, Fuller SD, Antony C, Krijnse-Locker J, Bartenschlager R. Composition and three-dimensional architecture of the dengue virus replication and assembly sites. Cell Host Microbe. 2009 Apr 23;5(4):365–75. doi: 10.1016/j.chom.2009.03.007.

43. Sangiambut S, Keelapang P, Aaskov J, Puttikhunt C, Kasinrerk W, Malasit P, Sittisombut N. Multiple regions in dengue virus capsid protein contribute to nuclear localization during virus infection. J Gen Virol. 2008 May;89(Pt 5):1254–1264. doi: 10.1099/vir.0.83264-0.

44. Zhang, X., Zhang, Y., Jia, R. et al. Structure and function of capsid protein in flavivirus infection and its applications in the development of vaccines and therapeutics. Vet Res 52, 98 (2021). https://doi.org/10.1186/s13567-021-00966-2

45. Mackenzie, J. M., Jones, M. K. & Young, P. R. Improved membrane preservation of flavivirus-infected cells with cryosectioning. Journal of Virological Methods 56, 67–75 (1996). https://doi.org/10.1016/0166-0934(95)01916-2.

46. Caldas, L.A., Azevedo, R.C., da Silva, J.L. et al. Microscopy analysis of Zika virus morphogenesis in mammalian cells. Sci Rep 10, 8370 (2020). https://doi.org/10.1038/s41598-020-65409-y

47. Westaway EG, Mackenzie JM, Kenney MT, Jones MK, Khromykh AA. Ultrastructure of Kunjin virus-infected cells: colocalization of NS1 and NS3 with double-stranded RNA, and of NS2B with NS3, in virus-induced membrane structures. J Virol. 1997 Sep;71(9):6650–61. doi: 10.1128/JVI.71.9.6650-6661.1997.

48. Aktepe TE, Mackenzie JM. Shaping the flavivirus replication complex: It is curvaceous! Cell Microbiol. 2018 Aug;20(8):e12884. doi: 10.1111/cmi.12884.

49. Mackenzie JM, Khromykh AA, Westaway EG. Stable expression of noncytopathic Kunjin replicons simulates both ultrastructural and biochemical characteristics observed during replication of Kunjin virus. Virology. 2001 Jan 5;279(1):161–72. doi: 10.1006/viro.2000.0691.

50. Hamel R, Dejarnac O, Wichit S, Ekchariyawat P, Neyret A, Luplertlop N, Perera-Lecoin M, Surasombatpattana P, Talignani L, Thomas F, Cao-Lormeau VM, Choumet V, Briant L, Desprès P, Amara A, Yssel H, Missé D. Biology of Zika Virus Infection in Human Skin Cells. J Virol. 2015 Sep;89(17):8880–96. doi: 10.1128/JVI.00354-15.

51. Barreto-Vieira DF, Jácome FC, da Silva MAN, Caldas GC, de Filippis AMB, de Sequeira PC, de Souza EM, Andrade AA, Manso PPA, Trindade GF, Lima SMB, Barth OM. Structural investigation of C6/36 and Vero cell cultures infected with a Brazilian Zika virus. PLoS One. 2017 Sep 12;12(9):e0184397. doi: 10.1371/journal.pone.0184397.

52. Stollar V, Schlesinger RW, Stevens TM. Studies on the nature of dengue viruses. III. RNA synthesis in cells infected with type 2dengue virus. Virology. 1967 Dec;33(4):650-8. doi: 10.1016/0042-6822(67)90065-7.

53. Ferreira GP, Trindade GS, Vilela JM, Da Silva MI, Andrade MS, Kroon EG. Climbing the steps of viral atomic force microscopy: visualization of Dengue virus particles. J Microsc. 2008 Jul;231(Pt 1):180–5. doi: 10.1111/j.1365-2818.2008.02028.x.

54. Sirohi D, Chen Z, Sun L, Klose T, Pierson TC, Rossmann MG, Kuhn RJ. The 3.8 Å resolution cryo-EM structure of Zika virus. Science. 2016 Apr 22;352(6284):467-70. doi: 10.1126/science.aaf5316.

55. Sevvana M, Long F, Miller AS, Klose T, Buda G, Sun L, Kuhn RJ, Rossmann MG. Refinement and Analysis of the Mature Zika Virus Cryo-EM Structure at 3.1 Å Resolution. Structure. 2018 Sep 4;26(9):1169–1177.e3. doi: 10.1016/j.str.2018.05.006.

56. Kostyuchenko VA, Chew PL, Ng TS, Lok SM. Near-atomic resolution cryo-electron microscopic structure of dengue serotype 4 virus. J Virol. 2014 Jan;88(1):477–82. doi: 10.1128/JVI.02641-13.

57. Lyonnais, S., Hénaut, M., Neyret, A. et al. Atomic force microscopy analysis of native infectious and inactivated SARS-CoV-2 virions. Sci Rep 11, 11885 (2021). https://doi.org/10.1038/s41598-021-91371-4

58. McGee CE, Shustov AV, Tsetsarkin K, Frolov IV, Mason PW, Vanlandingham DL, Higgs S. Infection, dissemination, and transmission of a West Nile virus green fluorescent protein infectious clone by Culex pipiens quinquefasciatus mosquitoes. Vector Borne Zoonotic Dis. 2010 Apr;10(3):267–74. doi: 10.1089/vbz.2009.0067.

59. Yin Z, Chen YL, Schul W, Wang QY, Gu F, Duraiswamy J, Kondreddi RR, Niyomrattanakit P, Lakshminarayana SB, Goh A, Xu HY, Liu W, Liu B, Lim JY, Ng CY, Qing M, Lim CC, Yip A, Wang G, Chan WL, Tan HP, Lin K, Zhang B, Zou G, Bernard KA, Garrett C, Beltz K, Dong M, Weaver M, He H, Pichota A, Dartois V, Keller TH, Shi PY. An adenosine nucleoside inhibitor of dengue virus. Proc Natl Acad Sci U S A. 2009 Dec 1;106(48):20435–9. doi: 10.1073/pnas.0907010106.

60. Deng YQ, Zhang NN, Li CF, Tian M, Hao JN, Xie XP, Shi PY, Qin CF. Adenosine Analog NITD008 Is a Potent Inhibitor of Zika Virus. Open Forum Infect Dis. 2016 Aug 30;3(4):ofw175. doi: 10.1093/ofid/ofw175.

61. Mishin VP, Cominelli F, Yamshchikov VF. A ’minimal’ approach in design of flavivirus infectious DNA. Virus Res. 2001 Dec 4;81(1-2):113–23. doi: 10.1016/s0168-1702(01)00371-9.

62. Santos JJ, Magalhães T, Silva Junior JV, Silva AN, Cordeiro MT, Gil LH. Full-length infectious clone of a low passage dengue virus serotype 2 from Brazil. Mem Inst Oswaldo Cruz. 2015 Aug;110(5):677–83. doi: 10.1590/0074-02760150053.

63. Hu M, Wu T, Yang Y, Chen T, Hao J, Wei Y, Luo T, Wu D, Li YP. Development and Characterization of a Genetically Stable Infectious Clone for a Genotype I Isolate of Dengue Virus Serotype 1. Viruses. 2022 Sep 18;14(9):2073. doi: 10.3390/v14092073.

64. Shan C, Xie X, Muruato AE, Rossi SL, Roundy CM, Azar SR, Yang Y, Tesh RB, Bourne N, Barrett AD, Vasilakis N, Weaver SC, Shi PY. An Infectious cDNA Clone of Zika Virus to Study Viral Virulence, Mosquito Transmission, and Antiviral Inhibitors. Cell Host Microbe. 2016 Jun 8;19(6):891–900. doi: 10.1016/j.chom.2016.05.004.

65. Santos JJ, Cordeiro MT, Bertani GR, Marques ET, Gil LH. A two-plasmid strategy for engineering a dengue virus type 3 infectious clone from primary Brazilian isolate. An Acad Bras Cienc. 2014 Dec;86(4):1749–59. doi: 10.1590/0001-3765201420130332.

66. Edmonds J, van Grinsven E, Prow N, Bosco-Lauth A, Brault AC, Bowen RA, Hall RA, Khromykh AA. A novel bacterium-free method for generation of flavivirus infectious DNA by circular polymerase extension reaction allows accurate recapitulation of viral heterogeneity. J Virol. 2013 Feb;87(4):2367–72. doi: 10.1128/JVI.03162-12.

67. Ávila-Pérez G, Nogales A, Martín V, Almazán F, Martínez-Sobrido L. Reverse Genetic Approaches for the Generation of Recombinant Zika Virus. Viruses. 2018 Oct 31;10(11):597. doi: 10.3390/v10110597.

68. Pu SY, Wu RH, Yang CC, Jao TM, Tsai MH, Wang JC, Lin HM, Chao YS, Yueh A. Successful propagation of flavivirus infectious cDNAs by a novel method to reduce the cryptic bacterial promoter activity of virus genomes. J Virol. 2011 Mar;85(6):2927–41. doi: 10.1128/JVI.01986-10.

69. Yun SI, Choi YJ, Yu XF, Song JY, Shin YH, Ju YR, Kim SY, Lee YM. Engineering the Japanese encephalitis virus RNA genome for the expression of foreign genes of various sizes: implications for packaging capacity and RNA replication efficiency. J Neurovirol. 2007 Dec;13(6):522–35. doi: 10.1080/13550280701684651.

70. Suphatrakul A, Duangchinda T, Jupatanakul N, Prasittisa K, Onnome S, Pengon J, Siridechadilok B. Multi-color fluorescent reporter dengue viruses with improved stability for analysis of a multi-virus infection. PLoS One. 2018 Mar 16;13(3):e0194399. doi: 10.1371/journal.pone.0194399.

71. Tan, T.Y., Fibriansah, G., Kostyuchenko, V.A. et al. Capsid protein structure in Zika virus reveals the flavivirus assembly process. Nat Commun 11, 895 (2020). https://doi.org/10.1038/s41467-020-14647-9

72. Barnard TR, Abram QH, Lin QF, Wang AB, Sagan SM. Molecular Determinants of Flavivirus Virion Assembly. Trends Biochem Sci. 2021 May;46(5):378–390. doi: 10.1016/j.tibs.2020.12.007.

73. Sirohi D, Chen Z, Sun L, Klose T, Pierson TC, Rossmann MG, Kuhn RJ. The 3.8 Å resolution cryo-EM structure of Zika virus. Science. 2016 Apr 22;352(6284):467-70. doi: 10.1126/science.aaf5316.

74. Kuhn RJ, Zhang W, Rossmann MG, Pletnev SV, Corver J, Lenches E, Jones CT, Mukhopadhyay S, Chipman PR, Strauss EG, Baker TS, Strauss JH. Structure of dengue virus: implications for flavivirus organization, maturation, and fusion. Cell. 2002 Mar 8;108(5):717–25. doi: 10.1016/s0092-8674(02)00660-8.

75. Zhang, W., Chipman, P., Corver, J. et al. Visualization of membrane protein domains by cryo-electron microscopy of dengue virus. Nat Struct Mol Biol 10, 907–912 (2003). https://doi.org/10.1038/nsb990

76. Therkelsen MD, Klose T, Vago F, Jiang W, Rossmann MG, Kuhn RJ. Flaviviruses have imperfect icosahedral symmetry. Proc Natl Acad Sci U S A. 2018 Nov 6;115(45):11608–11612. doi: 10.1073/pnas.1809304115.

77. Li XF, Li XD, Deng CL, Dong HL, Zhang QY, Ye Q, Ye HQ, Huang XY, Deng YQ, Zhang B, Qin CF. Visualization of a neurotropic flavivirus infection in mouse reveals unique viscerotropism controlled by host type I interferon signaling. Theranostics. 2017 Feb 12;7(4):912–925. doi: 10.7150/thno.16615.

78. Therkelsen MD, Klose T, Vago F, Jiang W, Rossmann MG, Kuhn RJ. Flaviviruses have imperfect icosahedral symmetry. Proc Natl Acad Sci U S A. 2018 Nov 6;115(45):11608–11612. doi: 10.1073/pnas.1809304115.

